# Optogenetic induction of chronic glucocorticoid exposure in early-life impairs stress-response in larval zebrafish

**DOI:** 10.1101/2022.09.09.507267

**Authors:** Jatin Nagpal, Helen Eachus, Olga Lityagina, Soojin Ryu

## Abstract

Organisms respond to stressors through a coordinated set of physiological and behavioural responses. Zebrafish provides an opportunity to study conserved mechanisms underlying the stress-response that is regulated largely by the neuroendocrine Hypothalamus-Pituitary-Adrenal/Interrenal (HPA) axis, with glucocorticoids (GC) as the final effector. In this study, we evaluated the effect of chronically active GC signalling in early life on the baseline and stress evoked GC(cortisol) levels in larval zebrafish. To this end, we employed an optogenetic actuator, *Beggiatoa* photoactivated adenylyl cyclase, expressed in the interrenal cells of zebrafish and demonstrate that its chronic activation leads to hypercortisolaemia and dampens the acute-stress evoked cortisol levels, across a variety of stressor modalities during early life. This blunting of stress-response, a phenotype reported by many studies to be observed in human subjects exposed to early-life trauma, was conserved in ontogeny at a later developmental stage. Furthermore, we observe a strong reduction of proopiomelanocortin (POMC)-expressing cells in the pituitary as well as global upregulation of FKBP5 gene expression, impinging on the negative feedback regulation elicited by elevated cortisol levels. Going forward, we propose that this model can be leveraged to tease apart the mechanisms underlying developmental programming of HPA axis by early-life stress and its implications for vulnerability and resilience to stress in adulthood.

## Introduction

Organisms respond to physical and psychological threats or stressors by mounting an integrated physiological and behavioural adaptive stress response, mediated by the neuroendocrine Hypothalamus-Pituitary-Adrenal (HPA) axis (Chrousos 2009). This allostatic maintenance of health can be challenged by cumulative chronic stressors, especially in the critical period of early-life, leading to compromised adaptive brain-body response and hence vulnerability to stress-related disorders including depression, anxiety and post-traumatic disorder (McEwen 2007; Lupien et al. 2009; Juster, McEwen, and Lupien 2010; van Bodegom, Homberg, and Henckens 2017).

The HPA axis is a cascade of physiological events culminating in the release of glucocorticoids (GC), which is the final effector of the stress response (de Kloet, Joëls, and Holsboer 2005). GC regulates its own release and terminates the stress response by negative feedback on the secretion of corticotropin-releasing hormone (CRH) and adrenocorticotropin hormone (ACTH) from the hypothalamus and proopiomelanocortin *(pomc)*-positive pituitary corticotrophs, respectively acting via the GC receptor (GR) (Charmandari, Tsigos, and Chrousos 2005). Another important modulator of the stress response is the co-chaperone FK506-binding protein 51 (FKBP5/FKBP51), a negative regulator of the GR. GR activation leads to the induction of FKBP5 expression, and with the inhibitory effect of FKBP5 on GR activity, it generates an ultra-short, negative feedback loop that regulates GR activity (Zannas et al. 2016). Additionally, polymorphisms in the FKBP5 gene have been associated with stress-related disorders in humans and rodents (Matosin, Halldorsdottir, and Binder 2018).

Early-life stress (ELS) led dysregulation of the HPA-axis can cause maladaptive developmental and functional maturation of stress response machinery, which may provide a mechanistic basis for altered stress susceptibility in later life (Taylor 2010; Maccari et al. 2014; Sandi and Haller 2015; Chen and Baram 2016). There is a vast literature covering effects of ELS on the structural and functional development of the HPA-axis and its key associated modulators (amygdala, hippocampus, and prefrontal cortex), particularly addressing neuroendocrine consequences (van Bodegom, Homberg, and Henckens 2017; Agorastos et al. 2019). These alterations induced by ELS, primarily studied in the rodent models, are dependent on the developmental time window affected, the sex of the individual, and the developmental stage at which effects are assessed. ELS-induced HPA axis dysfunction is a hallmark in a variety of neuropsychiatric diseases (e.g. depression and post-traumatic stress disorder, PTSD)(Pariante and Lightman 2008; Daskalakis et al. 2016). Many studies encompassing rodent and human subjects exposed to ELS or chronic stress have reported a blunted HPA axis response (Albeck et al. 1997; Heim et al. 2001; Burke et al. 2005; Cohen et al. 2006; Carpenter et al. 2007; Tomiyama, Dallman, and Epel 2011; Ouellet-Morin et al. 2011; Grimm et al. 2014; Carroll et al. 2017; Bunea, Szentágotai-Tătar, and Miu 2017; Lam et al. 2019; Xin et al. 2020). Also, exogenous treatment with GC or GR agonist have been shown to result in blunting of the endogenous stress response (Felszeghy, Bagdy, and Nyakas 2000; Kinlein, Wilson, and Karatsoreos 2015), indicating effects on GC negative feedback.

Larval zebrafish with their external development and excellent genetic and developmental analysis tools are emerging as an important complement to rodents in the study of stress on the brain and behaviour (Eachus, Choi, and Ryu 2021; de Abreu et al. 2021; Gerlai 2020). Importantly, the Hypothalamic-Pituitary-Interrenal (HPI) axis in zebrafish is homologous to the HPA axis in mammals (Wendelaar Bonga 1997; Flik et al. 2006) and larval zebrafish respond to stressors with increased cortisol (Alsop and Vijayan 2008; Alderman and Bernier 2009; Fuzzen et al. 2010; Steenbergen et al. 2011). Also, their basal cortisol levels and expression levels of genes involved in corticosteroid synthesis and signaling increase drastically around the time of hatching, indicative of a stress response that matures early in development (Alsop and Vijayan 2009; Alderman and Bernier 2009). Furthermore, transgenic approaches have identified evolutionarily conserved chemoarchitecture of hypothalamic and pituitary regions (Herget et al. 2014; Gutierrez-Triana et al. 2014; Nagpal et al. 2019) and methods have been established to manipulate the endogenous level of GCs using optogenetic tools thereby allowing controlled GC exposure with unprecedented specificity and temporal resolution (De Marco et al. 2013, 2016; Gutierrez-Triana et al. 2015). In this optogenetic approach, transgenic zebrafish with targeted expression of *beggiatoa* photoactivated adenylyl cyclase (bPAC) (Ryu et al. 2010; Stierl et al. 2011) both in pituitary corticotrophs, which produce ACTH (*pomc*:bPAC) (Rodrigo J. De Marco et al. 2013) and in steroidogenic cells that produce GCs within interrenal gland (*star*:bPAC) (Gutierrez-Triana et al. 2015) have been generated. Using this optogenetic approach, endogenous GC level was increased upon blue light stimulation, and it was demonstrated that the pituitary-adrenal system exerts a rapid organizing effect on behaviour in larval zebrafish (De Marco et al. 2016).

Here, we focus on investigating the basal and stress-induced cortisol changes upon chronic early-life HPA axis activation, using the optogenetically-induced non-invasive model of GC induction in larval zebrafish (Gutierrez-Triana et al. 2015). This mimic of early-life stress enabled us to profile cortisol levels at basal and upon acute-stress as a function of the chronically active HPA axis. We further investigated mechanisms embedded in the negative feedback of GC on POMC-expressing cells and FKBP5 in the model.

## Materials and Methods

### Zebrafish husbandry and maintenance

Zebrafish breeding and maintenance were performed under standard conditions on a 12:12 light/dark cycle (Westerfield 2000). *Tg(2kbStAR:bPAC-2A-tdTomato)hd19Tg* (Gutierrez-Triana et al. 2015) were bred with wild-type AB/TL zebrafish strain and the embryos were raised in egg water (3 g sea-salt/10 L ultrapure water) at 28 °C inside an incubator (RuMed 3101, Rubarth Apparate GmbH, Laatzen, Germany) with 12:12 light/dark cycle. Transgenic and non-transgenic siblings were screened at 3 days post fertilization (dpf) on a fluorescent stereomicroscope and transferred in petri dishes with egg water for experiments at later timepoints. Experiments were carried out with 6 dpf larvae. Larvae older than 6 dpf were transferred to plastic tanks with 400 ml of egg water in groups of thirty and fed with paramecia daily. Zebrafish experimental procedures were performed according to the guidelines of the German animal welfare law and approved by the local government (Regierungspräsidium Karlsruhe; G-29/12).

### Stressor treatment and cortisol assay

#### Chronic optogenetic induction of GC

Transgenic *Tg(2kbStAR:bPAC-2A-tdTomato)* and non-transgenic siblings were raised at 28 °C inside an incubator (RuMed 3101, Rubarth Apparate GmbH, Laatzen, Germany) with 12:12 light/dark cycle with light phase consisting of white light emitted by the standard white light lamp of the incubator. The continuous white light during the light phase of the circadian cycle was sufficient to activate the light-sensitive optogenetic protein bPAC expressed in the interrenal cells of the transgenic larvae, and thereby achieve chronic activation of the HPA axis from 0 dpf onwards. bPAC is activated strongly at UV-blue wavelengths of the spectrum (370 – 500 nm) with activation light intensities in the µW ranges (Ryu et al. 2010; Stierl et al. 2011; De Marco et al. 2013, 2016), which is amply provided by the white light illumination in the incubator. For experiments requiring avoiding unspecific activation of bPAC, *Tg(2kbStAR:bPAC-2A-tdTomato)* embryos were raised in custom-made containers covered by 550 nm long-pass filters (Thor-labs).

#### Osmotic and pH stressor

A hyperosmotic solution of 100 mM NaCl and low pH solution of 1mM HCl were employed as stressors (De Marco et al. 2013; Gutierrez-Triana et al. 2015; Castillo-Ramírez, Ryu, and De Marco 2019). Stock solutions of NaCl (from Sigma) and HCl (from Sigma) prepared in egg water were added to the petri dishes with zebrafish larvae (raised at fixed density for experiments) to achieve final concentrations of 100 mM NaCl and 1 mM HCl, which constituted the osmotic and pH stress, respectively. Samples were collected 10 minutes after the administration of stock solutions.

#### Mechanical stressor

A previously established mechanosensory stress protocol (Castillo-Ramírez, Ryu, and De Marco 2019) that used a magnetic bead in a dish placed on a magnetic plate, thereby generating unpredictable perturbation of the medium was employed. Petri dishes (inner diameter: 3.5 cm) containing 30 larvae and a plastic-covered stirrer (magnetic stir bar micro PTFE 6 mm x 3 mm; Fisher Scientific) were placed on a magnetic stir plate (Variomag Poly 15 stirrer plate; Thermo Scientific) and kept at 28 °C inside an incubator. The mechanosensory stress protocol was performed for 3 minutes at 330 rpm. Samples were collected 10 minutes after the onset of stimulation. Control larvae were collected after equal handling, omitting exposure to vortex flows (i.e., stir bars inside the Petri dishes were absent).

#### Whole body cortisol

The procedures for cortisol extraction were as previously described (De Marco et al. 2016). Unexposed larvae (control samples) were collected after equal handling, omitting stressor exposure. Larvae were immobilized using ice-cold water, frozen in an ethanol / dry ice bath and stored at –20°C for subsequent extraction. Each replicate consisted of 15 larvae and stimulation and sample collection were carried out between 10:30 and 13:00. The cortisol ELISA assay was performed as per manufacturer’s instructions using the Cisbio HTRF cortisol ELISA kit (62CRTPEG, Cisbio, Perkin Elmer) and measurements were performed on Tecan Infinite M1000 Pro multi-plate reader.

### Fluorescent in-situ hybridization

Fluorescent in-situ hybridization (ISH) was performed as described (Herget et al. 2014), using a *pomca* in situ probe (Herzog et al. 2003). Larvae were imaged in 80% glycerol in PBS using a 20x objective on a Leica SPE confocal microscope. Subsequent image processing and evaluation were performed on ImageJ as described (Herget et al. 2014).

### RT-qPCR

Larvae were snap frozen in dry ice, with each independent replicate comprising of 15 larvae. RNA was extracted using the Quick-RNA Microprep Kit (Zymo Research) after grinding the tissue with an electric homogenizer (Kimble, VWR). RNA concentration was measured using a Nanodrop ND2000. cDNA was prepared with the High-Capacity RNA-to-cDNA Kit (Applied Biosystems) using between 1.5 µg and 2 µg of total RNA per 20 µL-reaction. qPCR was performed in a 96-well plate (Sapphire Microplatte, Greiner) with PowerUp SYBR Green Mastermix (Applied Biosystems) in the CFX BioRad Real-Time PCR Detection System following its recommended protocols. Primers (fkbp5 F: 5′ GTGTTCGTCCACTACACC R: 5′ TCTCCTCACGATCCCACC; rpl13a F: 5′ TGGTGAGGTGTGAGGGTATCAAC R: 5′ AATTTGCGTGTGGGTTTCAGAC; sep15 F: 5′ TATTGTTGATTGTTGCTGAGGG R: 5′ ACGCTGAGAGATGTACACAGGA (Xu et al. 2016). For analysis, 2^−ΔΔCt^ method was applied using the rpl13a and sep15 as housekeeping genes for two independent experiments.

### Statistics

All data are shown as bars or single measurement points, mean and standard error of the mean (S.E.M.). ANOVA was used for multiple group comparisons followed by Tukey’s post hoc tests. Analyses were carried out using MS Excel (Microsoft; Redmond, WA, USA) and Prism 5 (Graphpad Software Inc.; San Diego, CA, USA).

## Results

### Specific and non-invasive activation of stress axis using optogenetics

To achieve chronic activation of the HPA(Interrenal) axis in a specific and non-invasive fashion, we utilized a previously reported (Gutierrez-Triana et al. 2015) transgenic line expressing the light-activated adenylyl cyclase bPAC in the steroioidogenic interrenal cells (Figure 1). This transgenic line employs a 2kb promoter element of the steroidogenic acute regulatory protein (StAR), a a rate-limiting mediator of steroid hormone biosynthesis (Stocco 2000), to drive the expression of Beggiatoa photoactivated adenylyl cyclase (bPAC) (Ryu et al. 2010; Stierl et al. 2011) coupled to the fluorescent protein TdTomato exclusively in larval steroidogenic interrenal cells. Upon binding to its receptor, ACTH leads to cAMP production and Ca^2+^ influx in adrenocortical cells, which triggers the synthesis and release of GC (Gallo-Payet and Payet 2003). Acute blue-light stimulation of the transgenic larvae has been shown to elicit transient hypercortisolemia (Gutierrez-Triana et al. 2015). bPAC, though with its peak activation in blue wavelength range, has a broad activation spectrum with high light sensitivity (in µW range)(Stierl et al. 2011; Ryu et al. 2010). We reasoned, to achieve chronic elevation of endogenous GC levels, the white light from the raising incubator in early-life (0 till 6 dpf) could be used to activate bPAC. Thus, the fish larvae were exposed to white light from 0 dpf to increase the cortisol output.

**Figure 1:**
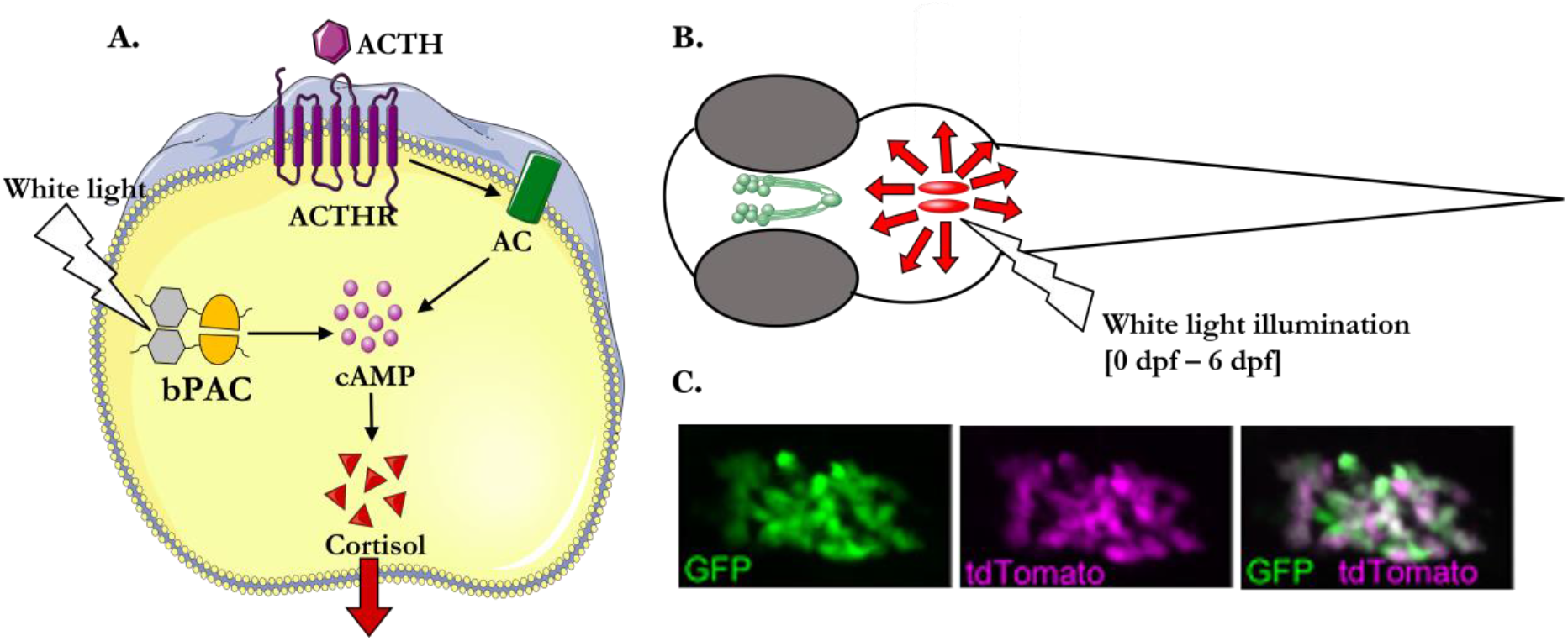
**A**. Schematic representation of the interrenal cells optogenetically activated using photoactivatable adenylyl cyclase bPAC, increasing cortisol output (dark red arrows). **B**. The hypothalamo-pituitary complex (green) causes the release of ACTH, which further acts on interrenal cells (red) to induce the release of cortisol (red arrows). Illumination with white light from 0 dpf onwards activates the interrenal cells, thereby increasing cortisol output. **C**. Colocalization of GFP and tdTomato in a double-transgenic 4-dpf larva expressing GFP and bPAC-2A-tdTomato under the control of the 2-kb star promoter, demonstrating the specificity of the *Tg(2kbStARp:bPAC-2A-tdTomato)* line (Gutierrez-Triana et al. 2015). Abbreviations: ACTHR ACTH receptor, AC Adenylyl cyclase, bPAC Beggiatoa photo-activated adenylyl cyclase.

### Chronic early life elevation of glucocorticoids leads to elevated basal cortisol levels and blunted stress response at 6 dpf and 12 dpf

As a function of chronic early-life HPA axis activation, we measured the basal and acute stress-evoked whole body cortisol levels of *Tg(2kbStARp:bPAC-2A-tdTomato)* larvae and non-transgenic siblings at 6 dpf. The basal cortisol levels were significantly elevated in transgenic fish as compared to the non-transgenic siblings, indicating a state of basal hypercortisolemia (one-way ANOVA p<0.01, followed by Tukey’s post hoc tests) (Figure 2A). The osmotic stress of 100 mM NaCl strongly elevated the cortisol levels in non-transgenic siblings indicating a physiological stress-response (one-way ANOVA p<0.001, followed by Tukey’s post hoc tests). However, there was no significant increase in the cortisol levels for the transgenic larvae (Figure 2A), demonstrating a blunted stress response. The similar phenotype of basal hypercortisolemia (one-way ANOVA p<0.001, followed by Tukey’s post hoc tests) and blunted cortisol response to osmotic stress was conserved in ontogeny at 12 dpf (Figure 2B).

**Figure 2.**
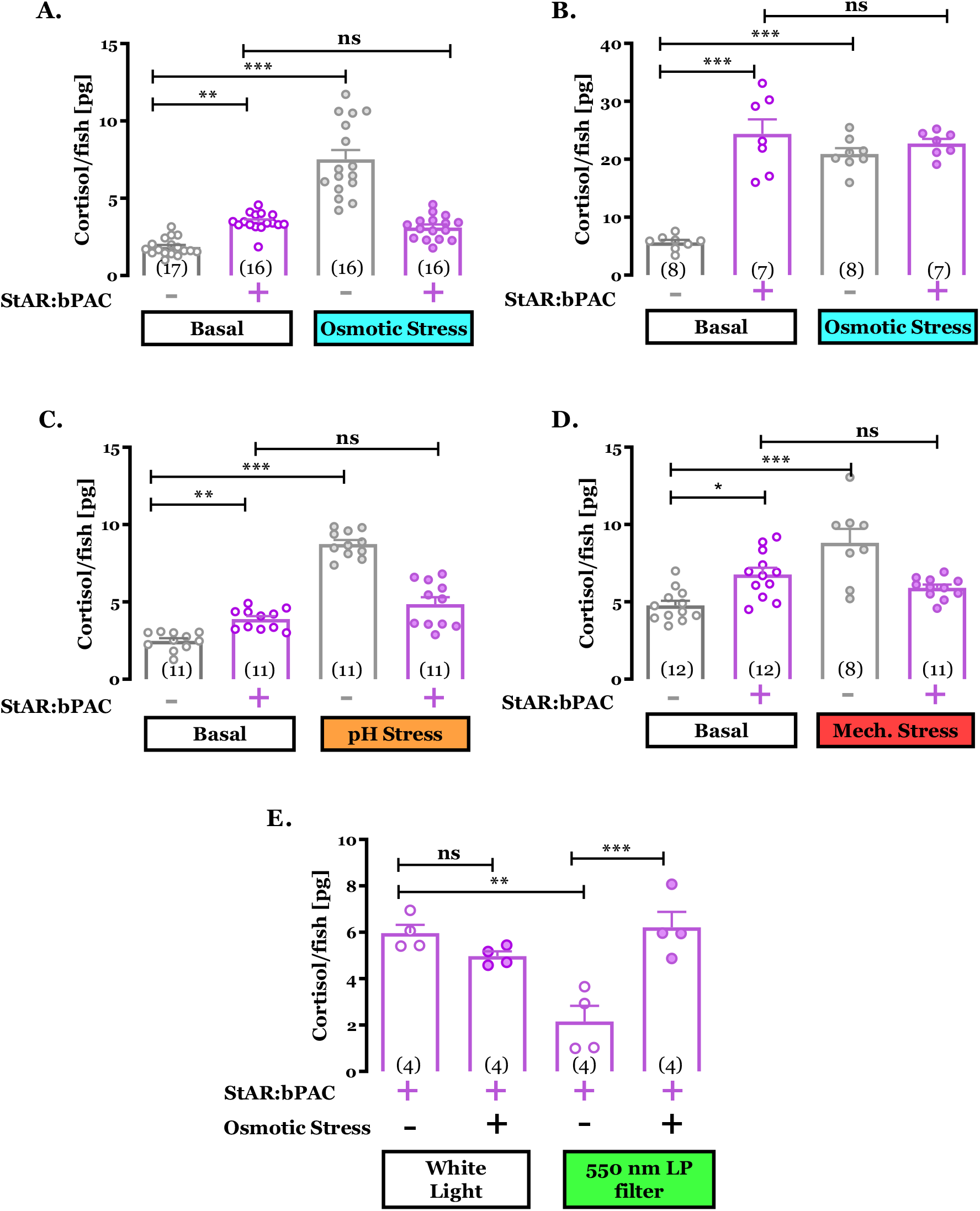
Chronic optogenetic induction of glucocorticoids leads to a state of basal hypercortisolemia and blunted cortisol response to stressors in *Tg(2kbStARp:bPAC-2A-tdTomato)* larvae relative to non-transgenic siblings. **A**. Basal (open circles) and 100mM NaCl osmotic stress (filled circles) evoked cortisol concentrations in bPAC-negative (−) (gray) and bPAC-positive (+) (purple) 6-dpf progenies of the Tg(2kbStARp:bPAC-2A-tdTomato) line. **B**. Basal and 100mM NaCl osmotic stress evoked cortisol concentrations in bPAC-negative (−) (gray) and bPAC-positive (+) (purple) 12-dpf progenies of the Tg(2kbStARp:bPAC-2A-tdTomato) line.. **C**. Basal and 1mM HCl pH stress evoked cortisol concentrations in bPAC-negative (−) (gray) and bPAC-positive (+) (purple) 6-dpf progenies of the Tg(2kbStARp:bPAC-2A-tdTomato) line **D**. Basal and mechanical stress evoked cortisol concentrations in bPAC-negative (−) (gray) and bPAC-positive (+) (purple) 6-dpf progenies of the Tg(2kbStARp:bPAC-2A-tdTomato) line. **E**. Basal and 100mM NaCl osmotic stress evoked cortisol concentrations in bPAC-positive (+) (purple) 6-dpf progenies of the Tg(2kbStARp:bPAC-2A-tdTomato) line, raised under activating white light and non-activating 550 nm long-pass (LP) filter. The number of samples (each comprising of 15 larvae) are mentioned in the parenthesis in individual bar plots for all conditions. The individual values are depicted in circles, and the bar represents mean with standard error of the mean. Statistical analyses involved one-way ANOVA followed by Tukey’s posthoc test: ns p > 0.05; * p ≤ 0.05; ** p ≤ 0.01; *** p ≤ 0.001.

We next wanted to investigate if the phenotype was dependent on stressor modality. To this end, we exposed the *Tg(2kbStARp:bPAC-2A-tdTomato)* transgenic larvae and non-transgenic siblings at 6 dpf to different stressors: pH stress (induced by 1mM HCl) and mechanical stress (induced by unpredictable perturbation of the medium by magnetic bead stirring) (Figure 2C, 2D). Significant basal hypercortisolemia was observed in transgenic larvae (one-way ANOVA p<0.05 and p<0.01, followed by Tukey’s post hoc tests), and importantly, the levels of cortisol were not elevated upon exposure to either of the stressors for transgenic larvae as compared to non-transgenic siblings. This indicated that the blunting of the cortisol response to acute stress was conserved for different stressor modalities.

### The blunted response is due to the hypercortisolic state induced by the activation of optogenetic protein bPAC

In order to ascertain that these response dynamics were causally linked with optogenetic activation of bPAC, we raised the *Tg(2kbStARp:bPAC-2A-tdTomato)* transgenic larvae under conditions (550 nm long pass filter) which restrict the activation light for bPAC and thus would inhibit the bPAC mediated optogenetic activation of interrenal cells (Figure 2E). We observed that the basal cortisol levels of filter-raised larvae at 6 dpf were significantly lower as compared to those larvae that had been raised under activating white light (one-way ANOVA p<0.01, followed by Tukey’s post hoc tests). Additionally, there was no blunting of the stress response observed as the there was a significant elevation of cortisol upon 100 mM NaCl exposure in filter-raised larvae (one-way ANOVA p<0.001, followed by Tukey’s post hoc tests).

### Effects of chronic GC induction on *pomc* and *fkbp5* gene expression

Having ascertained the effects on cortisol levels, we next wanted to explore the underlying mechanisms related to the negative feedback regulation by GC. We focused on two known targets of GC – *pomc* and *fkbp5* (Liu et al. 2003; Zannas et al. 2016). To evaluate the effect of chronic GC exposure on *pomc* expression, we performed Fluorescent ISH for the *pomc* gene (Herzog et al. 2003). At 6 dpf, we observed selective suppression of *pomc* expression in the anterior group of pituitary corticotrophs, known as the rostral pars distalis region, of *Tg(2kbStARp:bPAC-2A-tdTomato)* larvae relative to non-transgenic siblings at 6 dpf, suggesting a distinct domain that is responsive to glucocorticoid feedback (Figure 3A). Next, we performed a real-time quantitative PCR to evaluate the effects of chronic early life GC exposure on gene expression of *fkbp5*. The gene expression for *fkbp5* was strongly upregulated in *Tg(2kbStARp:bPAC-2A-tdTomato)* transgenic larvae compared to non-transgenic siblings raised in white light at 6 dpf (Figure 3B).

**Figure 3.**
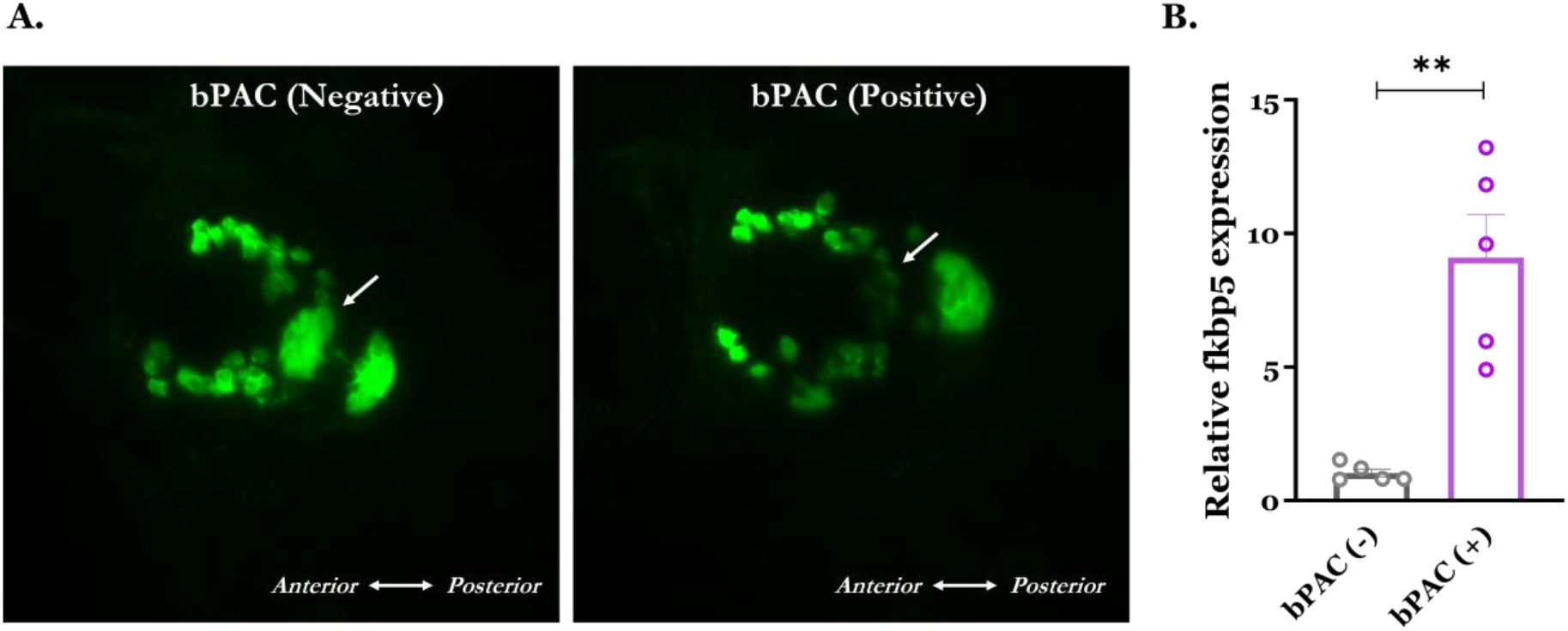
Effects of chronic early-life GC exposure on *pomc* and *fkbp5* gene expression A. Maximum intensity projection of confocal stacks show suppression of *pomc* (ISH staining) expression in the anterior domain of pituitary corticotrophs in *Tg(2kbStARp:bPAC-2A-tdTomato)* transgenic larvae relative to non-transgenic siblings at 6 dpf. B. RT-qPCR analysis for *fkbp5* expression shows strong organismal upregulation in *Tg(2kbStARp:bPAC-2A-tdTomato)* transgenic larvae relative to non-transgenic siblings at 6 dpf. Data is shown as mean with standard error of the mean, analysed with unpaired t-test (p-value<0.01). N=5 samples with 15 animals each.

## Discussion

Here, we have utilized an optogenetic approach to induce chronic HPA(I) axis activation through interrenal GC release in early-life in larval zebrafish. Specifically, we leveraged the broad activation spectrum (350-500nm) and high light sensitivity (µW range) of bPAC (Ryu et al. 2010; Stierl et al. 2011) to induce its activation in *Tg(2kbStARp:bPAC-2A-tdTomato)* transgenic fish (Gutierrez-Triana et al. 2015) via the white light used for raising the fish from 0 to 6 dpf. As a result of this bPAC-led optogenetic chronic activation of GC-producing interrenal cells, we observed significantly elevated levels of basal cortisol at 6 dpf, a developmental stage where the HPA(I) axis has matured in larval zebrafish and the larvae respond robustly to stressors with an increase in cortisol (Alsop and Vijayan 2008; De Marco et al. 2013). Importantly, cortisol is the final effector of stress response in zebrafish, as in humans, which is unlike the case in rodents, where corticosterone is the final mediator (Wendelaar Bonga 1997). The basal hypercortisolemia phenotype observed in this transgenic zebrafish achieved via direct, non-invasive and chronic activation of HPA axis, could serve as an endophenotype for human conditions displaying elevated baseline cortisol such as adrenal carcinoma or hyperplasia, cushing syndrome and neuropsychiatric conditions (e.g. depression) (Vreeburg et al. 2009; Dienes, Hazel, and Hammen 2013; Pariante 2016).

The other robust phenotype observed in our transgenic model across 3 different ecologically and ethologically relevant stressors (osmotic, pH, mechanical) is the severe blunting of the cortisol response to stress. This phenocopies the effects on the cortisol response observed in multiple studies where subjects (rodents and humans) were exposed to an early-life stress or trauma or GR agonist and their stress-responsivity was evaluated at a later time point (Ouellet-Morin et al. 2011; Kinlein, Wilson, and Karatsoreos 2015; Kinlein et al. 2019; Carpenter et al. 2007; Heim et al. 2001; Macmillan et al. 2008; Elzinga et al. 2008; Felszeghy, Bagdy, and Nyakas 2000). This endophenotypic conservation with models and cases of early-life trauma and post-traumatic stress disorder (PTSD) lends this transgenic zebrafish model to be used in the future to delve deeper into the aetiology and underlying mechanisms. Importantly, both phenotypes of basal hypercortisolaemia and blunted cortisol response to acute stress were not observed in bPAC restrictive illumination conditions (raising the larvae under 550 nm LP filter), providing evidence of the optogenetic interrenal activation as the underlying cause for the cortisol observations.

To understand and validate the underlying physiology underlying these chronic GC exposure-mediated phenotypes, we focused on two well-studied targets of GC, particularly in the context of the negative feedback loops of GC regulation – *pomc-*expressing pituitary corticotrophs and GR co-chaperone FKBP5. We observed strong and selective suppression of *pomc* expression in the rostral pars distalis of the pituitary gland in transgenic larval zebrafish at 6 dpf. This is similar to reports wherein larval zebrafish were exogenously treated with dexamethasone, a selective GR agonist (Liu et al. 2003; To et al. 2007; Peles, Swaminathan, and Levkowitz 2022), demonstrating the effect of greater GC negative feedback control and that this group of pituitary *pomc*-expressing cells is involved in the control of interrenal steroidogenesis and cortisol synthesis. This suppression of *pomc* expression thus could be a contributing factor for the blunted GC phenotype owing possibly due to lack of ACTH release upon stressor exposure. FKBP5 is a co-chaperone in the steroid receptor complex, whose expression is rapidly induced by GR activation. It exerts an inhibitory effect on intracellular GC signalling, thereby constituting an ultra-short negative feedback control that regulates GR activity and shapes neuroendocrine and stress reactivity (Matosin, Halldorsdottir, and Binder 2018; Häusl et al. 2019). There is also strong support in the literature for its role in shaping HPA axis function and overall stress reactivity (Touma et al. 2011; Häusl et al. 2021; Brix et al. 2022). Several human and animal studies have implicated interactions between stress and FKBP5 genotype on psychiatric conditions (e.g. PTSD, depression, anxiety) and related endophenotypes (Zannas et al. 2016). Our observation of strong organismal up-regulation of fkbp5 gene expression at 6 dpf following chronic GC exposure mirrors previous findings of dramatic induction of FKBP5 expression in a number of brain regions after stimulation with dexamethasone or stress exposure(Scharf et al. 2011). This overall mimic of FKBP5 disinhibition in our transgenic model may be of translational relevance since fkbp5 is a promising drug target for a variety of psychiatric and other disorders (Schmidt et al. 2012).

We can envisage that this zebrafish model will be of great relevance to look mechanistically at the developmental programming of HPA axis as well as adult stress-related and social behaviour, particularly as a function of early-life stress (Nagpal et al. 2019; Eachus, Choi, and Ryu 2021). Specifically, the genetic, developmental and high-throughput nature of the zebrafish system coupled to this model could be leveraged to reveal the molecular and cellular underpinnings of GC-mediated effects on brain, behaviour and organismal physiology (Griffiths et al. 2012; van den Bos et al. 2020; Lopez et al. 2021; Swaminathan et al. 2022). Furthermore, relatively understudied brain-body communication factors such as microbiome could be an important piece of the puzzle of GC – brain – behaviour axis, given that microbiome has been shown to strongly impact HPA axis function and stress-social behaviour (Nagpal and Cryan 2021). Advances in optogenetic tool development, particularly tools that have red-shifted spectral activation and low dark-activity could provide improved spatio-temporal control of GC levels and function (Fomicheva et al. 2019; Yang et al. 2021; Henß et al. 2021; Lehtinen, Nokia, and Takala 2021).

## Acknowledgements and Funding

SR is supported by the Mireille Gillings Foundation. SR is also supported by the German Federal Office for Education and Research grant number 01GQ1404. SR received support from DFG SPP1926 ‘Next generation optogenetics’. JN would like to acknowledge support from University Medical Center, Johannes Gutenberg University Mainz (Intramural Research Funding, Stufe1) and Irish Research Council (GOIPD/2019/714). JN would also like to thank Prof. John F. Cryan for his mentorship and Dr. María Rodriguez Aburto for her support. We would like to acknowledge the support of Ms. Kathrin Domdera for expert zebrafish maintenance, IMB Mainz Imaging facility for microscopy and Dr. Min-Kyeung Choi for comments on the manuscript.

## Author Contributions

Jatin Nagpal: Experiments were designed, performed and analyzed, funding was provided, the manuscript was written. Helen Eachus: Experiments were designed, performed and analyzed, the manuscript was written. Olga Lityagina: Experiments were performed and analyzed. Soojin Ryu: Experiments were designed, funding was provided, the manuscript was written.

## References

Abreu, Murilo S. de, Konstantin A. Demin, Ana C. V. V. Giacomini, Tamara G. Amstislavskaya, Tatyana Strekalova, Gleb O. Maslov, Yury Kositsin, Elena V. Petersen, and Allan V. Kalueff. 2021. “Understanding How Stress Responses and Stress-Related Behaviors Have Evolved in Zebrafish and Mammals.” Neurobiology of Stress 15 (100405): 100405.

Agorastos, Agorastos, Panagiota Pervanidou, George P. Chrousos, and Dewleen G. Baker. 2019. “Developmental Trajectories of Early Life Stress and Trauma: A Narrative Review on Neurobiological Aspects Beyond Stress System Dysregulation.” Frontiers in Psychiatry / Frontiers Research Foundation 10 (March): 118.

Albeck, McKittrick, Blanchard, Blanchard, Nikulina, McEwen, and Sakai. 1997. “Chronic Social Stress Alters Levels of Corticotropin-Releasing Factor and Arginine Vasopressin MRNA in Rat Brain.” The Journal of Neuroscience: The Official Journal of the Society for Neuroscience. http://eutils.ncbi.nlm.nih.gov/entrez/eutils/elink.fcgi?dbfrom=pubmed&id=9169547&retmode=ref&cmd=prlinks.

Alsop, Derek, and Mathilakath M. Vijayan. 2008. “Development of the Corticosteroid Stress Axis and Receptor Expression in Zebrafish.” American Journal of Physiology. Regulatory, Integrative and Comparative Physiology 294 (3): R711–9.

Bodegom, Miranda van, Judith R. Homberg, and Marloes J. A. G. Henckens. 2017. “Modulation of the Hypothalamic-Pituitary-Adrenal Axis by Early Life Stress Exposure.” Frontiers in Cellular Neuroscience 11 (April): 87.

Bos, Ruud van den, Suzanne Cromwijk, Katharina Tschigg, Joep Althuizen, Jan Zethof, Robert Whelan, Gert Flik, and Marcel Schaaf. 2020. “Early Life Glucocorticoid Exposure Modulates Immune Function in Zebrafish (Danio Rerio) Larvae.” Frontiers in Immunology 11 (April): 727.

Brix, Lea M., Alexander S. Häusl, Irmak Toksöz, Joeri Bordes, Lotte van Doeselaar, Clara Engelhardt, Sowmya Narayan, et al. 2022. “The Co-Chaperone FKBP51 Modulates HPA Axis Activity and Age-Related Maladaptation of the Stress System in Pituitary Proopiomelanocortin Cells.” Psychoneuroendocrinology 138 (April): 105670.

Bunea, Ioana Maria, Aurora Szentágotai-Tătar, and Andrei C. Miu. 2017. “Early-Life Adversity and Cortisol Response to Social Stress: A Meta-Analysis.” Translational Psychiatry 7 (12): 1274.

Burke, Heather M., Lia C. Fernald, Paul J. Gertler, and Nancy E. Adler. 2005. “Depressive Symptoms Are Associated With Blunted Cortisol Stress Responses in Very Low-Income Women.” Psychosomatic Medicine 67 (2): 211–16.

Carpenter, Linda L., John P. Carvalho, Audrey R. Tyrka, Lauren M. Wier, Andrea F. Mello, Marcelo F. Mello, George M. Anderson, Charles W. Wilkinson, and Lawrence H. Price. 2007. “Decreased Adrenocorticotropic Hormone and Cortisol Responses to Stress in Healthy Adults Reporting Significant Childhood Maltreatment.” Biological Psychiatry 62 (10): 1080–87.

Carroll, Douglas, Annie T. Ginty, Anna C. Whittaker, William R. Lovallo, and Susanne R. de Rooij. 2017. “The Behavioural, Cognitive, and Neural Corollaries of Blunted Cardiovascular and Cortisol Reactions to Acute Psychological Stress.” Neuroscience and Biobehavioral Reviews 77 (June): 74–86.

Castillo-Ramírez, Luis A., Soojin Ryu, and Rodrigo J. De Marco. 2019. “Active Behaviour during Early Development Shapes Glucocorticoid Reactivity.” Scientific Reports 9 (1): 12796.

Charmandari, Evangelia, Constantine Tsigos, and George Chrousos. 2005. “Endocrinology of the Stress Response.” Annual Review of Physiology 67 (1): 259–84.

Chen, Yuncai, and Tallie Z. Baram. 2016. “Toward Understanding How Early-Life Stress Reprograms Cognitive and Emotional Brain Networks.” Neuropsychopharmacology: Official Publication of the American College of Neuropsychopharmacology 41 (1): 197–206.

Chrousos, George P. 2009. “Stress and Disorders of the Stress System.” Nature Reviews. Endocrinology 5 (7): 374–81.

Cohen, Hagit, Joseph Zohar, Yori Gidron, Michael A. Matar, Dana Belkind, Uri Loewenthal, Nitsan Kozlovsky, and Zeev Kaplan. 2006. “Blunted HPA Axis Response to Stress Influences Susceptibility to Posttraumatic Stress Response in Rats.” Biological Psychiatry 59 (12): 1208–18.

Daskalakis, Nikolaos P., Hagit Cohen, Caroline M. Nievergelt, Dewleen G. Baker, Joseph D. Buxbaum, Scott J. Russo, and Rachel Yehuda. 2016. “New Translational Perspectives for Blood-Based Biomarkers of PTSD: From Glucocorticoid to Immune Mediators of Stress Susceptibility.” Experimental Neurology 284: 133–40.

De Marco, Rodrigo J., Antonia H. Groneberg, Chen-Min Yeh, Luis A. Castillo Ramírez, and Soojin Ryu. 2013. “Optogenetic Elevation of Endogenous Glucocorticoid Level in Larval Zebrafish.” Frontiers in Neural Circuits 7 (May): 82.

De Marco, Rodrigo J., Theresa Thiemann, Antonia H. Groneberg, Ulrich Herget, and Soojin Ryu. 2016. “Optogenetically Enhanced Pituitary Corticotroph Cell Activity Post-Stress Onset Causes Rapid Organizing Effects on Behaviour.” Nature Communications 7 (September): 12620.

Dienes, Kimberly A., Nicholas A. Hazel, and Constance L. Hammen. 2013. “Cortisol Secretion in Depressed, and at-Risk Adults.” Psychoneuroendocrinology 38 (6): 927–40.

Eachus, Helen, Min-Kyeung Choi, and Soojin Ryu. 2021. “The Effects of Early Life Stress on the Brain and Behaviour: Insights from Zebrafish Models.” Frontiers in Cell and Developmental Biology 9 (July): 657591.

Elzinga, Bernet M., Karin Roelofs, Marieke S. Tollenaar, Patricia Bakvis, Johannes van Pelt, and Philip Spinhoven. 2008. “Diminished Cortisol Responses to Psychosocial Stress Associated with Lifetime Adverse Events a Study among Healthy Young Subjects.” Psychoneuroendocrinology 33 (2): 227–37.

Felszeghy, K., G. Bagdy, and C. Nyakas. 2000. “Blunted Pituitary-Adrenocortical Stress Response in Adult Rats Following Neonatal Dexamethasone Treatment.” Journal of Neuroendocrinology 12 (10): 1014–21.

Fomicheva, Anastasia, Chen Zhou, Qian-Quan Sun, and Mark Gomelsky. 2019. “Engineering Adenylate Cyclase Activated by Near-Infrared Window Light for Mammalian Optogenetic Applications.” ACS Synthetic Biology 8 (6): 1314–24.

Gallo-Payet, Nicole, and Marcel D. Payet. 2003. “Mechanism of Action of ACTH: Beyond CAMP.” Microscopy Research and Technique 61 (3): 275–87.

Gerlai, Robert. 2020. “Evolutionary Conservation, Translational Relevance and Cognitive Function: The Future of Zebrafish in Behavioral Neuroscience.” Neuroscience and Biobehavioral Reviews 116 (September): 426–35.

Griffiths, Brian B., Peter J. Schoonheim, Limor Ziv, Lisa Voelker, Herwig Baier, and Ethan Gahtan. 2012. “A Zebrafish Model of Glucocorticoid Resistance Shows Serotonergic Modulation of the Stress Response.” Frontiers in Behavioral Neuroscience 6 (October): 1–10.

Grimm, Simone, Karin Pestke, Melanie Feeser, Sabine Aust, Anne Weigand, Jue Wang, Katja Wingenfeld, et al. 2014. “Early Life Stress Modulates Oxytocin Effects on Limbic System during Acute Psychosocial Stress.” Social Cognitive and Affective Neuroscience 9 (11): 1828–35.

Gutierrez-Triana, Jose Arturo, Ulrich Herget, Luis A. Castillo-Ramirez, Markus Lutz, Chen Min Yeh, Rodrigo J. De Marco, and Soojin Ryu. 2015. “Manipulation of Interrenal Cell Function in Developing Zebrafish Using Genetically Targeted Ablation and an Optogenetic Tool.” Endocrinology 156 (9): 3394–3401.

Gutierrez-Triana, Jose Arturo, Ulrich Herget, Patrick Lichtner, Luis A. Castillo-Ramírez, and Soojin Ryu. 2014. “A Vertebrate-Conserved Cis-Regulatory Module for Targeted Expression in the Main Hypothalamic Regulatory Region for the Stress Response.” BMC Developmental Biology 14 (1): 41.

Häusl, Alexander S., Georgia Balsevich, Nils C. Gassen, and Mathias V. Schmidt. 2019. “Focus on FKBP51: A Molecular Link between Stress and Metabolic Disorders.” Molecular Metabolism 29 (November): 170–81.

Häusl, Alexander S., Lea M. Brix, Jakob Hartmann, Max L. Pöhlmann, Juan-Pablo Lopez, Danusa Menegaz, Elena Brivio, et al. 2021. “The Co-Chaperone Fkbp5 Shapes the Acute Stress Response in the Paraventricular Nucleus of the Hypothalamus of Male Mice.” Molecular Psychiatry 26 (7): 3060–76.

Heim, C., D. J. Newport, R. Bonsall, A. H. Miller, and C. B. Nemeroff. 2001. “Altered Pituitary-Adrenal Axis Responses to Provocative Challenge Tests in Adult Survivors of Childhood Abuse.” The American Journal of Psychiatry 158 (4): 575–81.

Henß, Thilo, Jatin Nagpal, Shiqiang Gao, Ulrike Scheib, Alessia Pieragnolo, Alexander Hirschhäuser, Franziska Schneider-Warme, Peter Hegemann, Georg Nagel, and Alexander Gottschalk. 2021. “Optogenetic Tools for Manipulation of Cyclic Nucleotides Functionally Coupled to Cyclic Nucleotide-Gated Channels.” British Journal of Pharmacology, no. bph.15445 (March). https://doi.org/10.1111/bph.15445.

Herget, Ulrich, Andrea Wolf, Mario F. Wullimann, and Soojin Ryu. 2014. “Molecular Neuroanatomy and Chemoarchitecture of the Neurosecretory Preoptic-Hypothalamic Area in Zebrafish Larvae.” The Journal of Comparative Neurology 522 (7): 1542–64.

Herzog, Wiebke, Xianchun Zeng, Zsolt Lele, Carmen Sonntag, Jing Wen Ting, Chi Yao Chang, and Matthias Hammerschmidt. 2003. “Adenohypophysis Formation in the Zebrafish and Its Dependence on Sonic Hedgehog.” Developmental Biology 254 (1): 36–49.

Juster, Robert-Paul, Bruce S. McEwen, and Sonia J. Lupien. 2010. “Allostatic Load Biomarkers of Chronic Stress and Impact on Health and Cognition.” Neuroscience and Biobehavioral Reviews 35 (1): 2–16.

Kinlein, Scott A., Derrick J. Phillips, Chandler R. Keller, and Ilia N. Karatsoreos. 2019. “Role of Corticosterone in Altered Neurobehavioral Responses to Acute Stress in a Model of Compromised Hypothalamic-Pituitary-Adrenal Axis Function.” Psychoneuroendocrinology 102 (April): 248–55.

Kinlein, Scott A., Christopher D. Wilson, and Ilia N. Karatsoreos. 2015. “Dysregulated Hypothalamic-Pituitary-Adrenal Axis Function Contributes to Altered Endocrine and Neurobehavioral Responses to Acute Stress.” Frontiers in Psychiatry / Frontiers Research Foundation 6 (March): 31.

Kloet, E. Ron de, Marian Joëls, and Florian Holsboer. 2005. “Stress and the Brain: From Adaptation to Disease.” Nature Reviews. Neuroscience 6 (6): 463–75.

Lam, Jovian C. W., Grant S. Shields, Brian C. Trainor, George M. Slavich, and Andrew P. Yonelinas. 2019. “Greater Lifetime Stress Exposure Predicts Blunted Cortisol but Heightened DHEA Responses to Acute Stress.” Stress and Health: Journal of the International Society for the Investigation of Stress 35 (1): 15–26.

Lehtinen, Kimmo, Miriam S. Nokia, and Heikki Takala. 2021. “Red Light Optogenetics in Neuroscience.” Frontiers in Cellular Neuroscience 15: 778900.

Liu, Ning-Ai, Haigen Huang, Zhongan Yang, Wiebke Herzog, Matthias Hammerschmidt, Shuo Lin, and Shlomo Melmed. 2003. “Pituitary Corticotroph Ontogeny and Regulation in Transgenic Zebrafish.” Molecular Endocrinology 17 (5): 959–66.

Lopez, Juan Pablo, Elena Brivio, Alice Santambrogio, Carlo De Donno, Aron Kos, Miriam Peters, Nicolas Rost, et al. 2021. “Single-Cell Molecular Profiling of All Three Components of the HPA Axis Reveals Adrenal ABCB1 as a Regulator of Stress Adaptation.” Science Advances 7 (5). https://doi.org/10.1126/sciadv.abe4497.

Lupien, Sonia J., Bruce S. McEwen, Megan R. Gunnar, and Christine Heim. 2009. “Effects of Stress throughout the Lifespan on the Brain, Behaviour and Cognition.” Nature Reviews Neuroscience. https://doi.org/10.1038/nrn2639.

Maccari, S., H. J. Krugers, S. Morley-Fletcher, M. Szyf, and P. J. Brunton. 2014. “The Consequences of Early-Life Adversity: Neurobiological, Behavioural and Epigenetic Adaptations.” Journal of Neuroendocrinology 26 (10): 707–23.

Macmillan, Harriet L., Katholiki Georgiades, Eric K. Duku, Alison Shea, Meir Steiner, Anne Niec, Masako Tanaka, et al. 2008. “Cortisol Response to Stress in Female Youths Exposed to Childhood Maltreatment : Results of the Youth Mood Project.” BPS 66 (1): 62–68.

Matosin, Natalie, Thorhildur Halldorsdottir, and Elisabeth B. Binder. 2018. “Understanding the Molecular Mechanisms Underpinning Gene by Environment Interactions in Psychiatric Disorders: The FKBP5 Model.” Biological Psychiatry, no. 8: 1–10.

McEwen, Bruce S. 2007. “Physiology and Neurobiology of Stress and Adaptation: Central Role of the Brain.” Physiological Reviews 87: 873–904.

Nagpal, Jatin, and John F. Cryan. 2021. “Microbiota-Brain Interactions: Moving toward Mechanisms in Model Organisms.” Neuron 109 (24): 3930–53.

Nagpal, Jatin, Ulrich Herget, Min K. Choi, and Soojin Ryu. 2019. “Anatomy, Development, and Plasticity of the Neurosecretory Hypothalamus in Zebrafish.” Cell and Tissue Research 375 (1): 5–22.

Ouellet-Morin, Isabelle, Candice L. Odgers, Andrea Danese, Lucy Bowes, Sania Shakoor, Andrew S. Papadopoulos, Avshalom Caspi, Terrie E. Moffitt, and Louise Arseneault. 2011. “Blunted Cortisol Responses to Stress Signal Social and Behavioral Problems among Maltreated/Bullied 12-Year-Old Children.” Biological Psychiatry 70 (11): 1016–23.

Pariante, Carmine M. 2016. “The Glucocorticoid Receptor: Part of the Solution or Part of the Problem?:” Journal of Psychopharmacology. SAGE PublicationsLondon, Thousand Oaks, CA and New Delhi. http://journals.sagepub.com/doi/10.1177/1359786806066063.

Pariante, Carmine M., and Stafford L. Lightman. 2008. “The HPA Axis in Major Depression: Classical Theories and New Developments.” Trends in Neurosciences. Elsevier Current Trends. http://linkinghub.elsevier.com/retrieve/pii/S0166223608001641.

Peles, Guy, Amrutha Swaminathan, and Gil Levkowitz. 2022. “Glucocorticoid-Sensitive Period of Corticotroph Development – Implications in Early Life Stress.” BioRxiv. https://doi.org/10.1101/2022.06.13.495881.

Ryu, Min-Hyung, Oleg V. Moskvin, Jessica Siltberg-Liberles, and Mark Gomelsky. 2010. “Natural and Engineered Photoactivated Nucleotidyl Cyclases for Optogenetic Applications.” The Journal of Biological Chemistry 285 (53): 41501–8.

Sandi, Carmen, and József Haller. 2015. “Stress and the Social Brain: Behavioural Effects and Neurobiological Mechanisms.” Nature Reviews. Neuroscience 16 (5): 290–304.

Scharf, Sebastian H., Claudia Liebl, Elisabeth B. Binder, Mathias V. Schmidt, and Marianne B. Müller. 2011. “Expression and Regulation of the Fkbp5 Gene in the Adult Mouse Brain.” PloS One 6 (2): e16883.

Schmidt, Mathias V., Marcelo Paez-Pereda, Florian Holsboer, and Felix Hausch. 2012. “The Prospect of FKBP51 as a Drug Target.” ChemMedChem 7 (8): 1351–59.

Stierl, Manuela, Patrick Stumpf, Daniel Udwari, Ronnie Gueta, Rolf Hagedorn, Aba Losi, Wolfgang Gärtner, et al. 2011. “Light Modulation of Cellular CAMP by a Small Bacterial Photoactivated Adenylyl Cyclase, BPAC, of the Soil Bacterium Beggiatoa.” The Journal of Biological Chemistry 286 (2): 1181–88.

Stocco, D. M. 2000. “The Role of the StAR Protein in Steroidogenesis: Challenges for the Future.” The Journal of Endocrinology 164 (3): 247–53.

Swaminathan, Amrutha, Michael Gliksberg, Savani Anbalagan, Noa Wigoda, and Gil Levkowitz. 2022. “Stress Resilience Is Established during Development and Is Regulated by Complement Factors.” BioRxiv. https://doi.org/10.1101/2022.01.31.478444.

Taylor, Shelley E. 2010. “Mechanisms Linking Early Life Stress to Adult Health Outcomes.” Proceedings of the National Academy of Sciences of the United States of America 107 (19): 8507–12.

To, Thuy Thanh, Stefanie Hahner, Gabriela Nica, Klaus B. Rohr, Matthias Hammerschmidt, Christoph Winkler, and Bruno Allolio. 2007. “Pituitary-Interrenal Interaction in Zebrafish Interrenal Organ Development.” Molecular Endocrinology 21 (2): 472–85.

Tomiyama, A. Janet, Mary F. Dallman, and Elissa S. Epel. 2011. “Comfort Food Is Comforting to Those Most Stressed: Evidence of the Chronic Stress Response Network in High Stress Women.” Psychoneuroendocrinology 36 (10): 1513–19.

Touma, Chadi, Nils Christian Gassen, Leonie Herrmann, Joyce Cheung-Flynn, Dominik R. Büll, Irina A. Ionescu, Jan-Michael Heinzmann, et al. 2011. “FK506 Binding Protein 5 Shapes Stress Responsiveness: Modulation of Neuroendocrine Reactivity and Coping Behavior.” Biological Psychiatry 70 (10): 928–36.

Vreeburg, Sophie A., ;. Witte, J. G. Hoogendijk, ;. Johannes Van Pelt, Roel H. Derijk, Jolanda C. M. Verhagen, Bsc ;. Richard Van Dyck, Johannes H. Smit, Frans G. Zitman, and Brenda W. J. H. Penninx. 2009. “Major Depressive Disorder and Hypothalamic-Pituitary-Adrenal Axis Activity Results From a Large Cohort Study.” Vol. 66.

Wendelaar Bonga, S. E. 1997. “The Stress Response in Fish.” Physiological Reviews 77 (3): 591–625.

Westerfield, M. 2000. “The Zebrafish Book : A Guide for the Laboratory Use of Zebrafish.” http://Zfin.Org/Zf_info/Zfbook/Zfbk.Html. https://ci.nii.ac.jp/naid/10029409142/.

Xin, Yuanyuan, Zhuxi Yao, Weiwen Wang, Yuejia Luo, André Aleman, and Jianhui Wu. 2020. “Recent Life Stress Predicts Blunted Acute Stress Response and the Role of Executive Control.” Stress 23 (3): 359–67.

Xu, H., C. Li, Q. Zeng, I. Agrawal, X. Zhu, and Z. Gong. 2016. “Genome-Wide Identification of Suitable Zebrafish Danio Rerio Reference Genes for Normalization of Gene Expression Data by RT-QPCR.” Journal of Fish Biology 88 (6): 2095–2110.

Yang, Shang, Oana M. Constantin, Divya Sachidanandan, Hannes Hofmann, Tobias C. Kunz, Vera Kozjak-Pavlovic, Thomas G. Oertner, et al. 2021. “PACmn for Improved Optogenetic Control of Intracellular CAMP.” BMC Biology 19 (1): 227.

Zannas, Anthony S., Tobias Wiechmann, Nils C. Gassen, and Elisabeth B. Binder. 2016. “Gene-Stress-Epigenetic Regulation of FKBP5: Clinical and Translational Implications.” Neuropsychopharmacology: Official Publication of the American College of Neuropsychopharmacology 41 (1): 261–74.

